# Disease-Linked Super-Trafficking of a Mutant Potassium Channel

**DOI:** 10.1101/2020.07.29.227231

**Authors:** Hui Huang, Laura M. Chamness, Carlos G. Vanoye, Georg Kuenze, Jens Meiler, Alfred L. George, Jonathan P. Schlebach, Charles R. Sanders

**Author notes:** **Corresponding author:** Charles R. Sanders. **One Sentence Summary:** Super-trafficking of R231C KCNQ1 was discovered and characterized as a new paradigm for protein trafficking/disease relationships.

## Abstract

Gain-of-function (GOF) mutations in the KCNQ1 voltage-gated potassium channel can induce cardiac arrhythmia. We tested whether any of the known GOF disease mutations in KCNQ1 act by increasing the amount of KCNQ1 that reaches the cell surface—“super-trafficking”. We found that levels of R231C KCNQ1 in the plasma membrane are 5-fold higher than wild type KCNQ1. This arises from both enhanced translocon-mediated membrane integration of the S4 voltage-sensor helix and an energetic linkage of C231 with the V129 and F166 side chains. Whole-cell electrophysiology recordings confirmed that R231C KCNQ1 in complex with KCNE1 is constitutively active, but also revealed the single channel activity of this mutant to be only 20% that of WT. The GOF phenotype associated with R231C therefore reflects the net effects of super-trafficking, reduced single channel activity, and constitutive channel activation. These investigations document membrane protein super-trafficking as a contributing mechanism to human disease.

## INTRODUCTON

The KCNQ1 (K_V_7.1) voltage-gated potassium channel plays a critical role in the cardiac action potential (*1–3*). Inherited missense mutations in *KCNQ1* encode amino acid changes in the protein that can trigger either loss of channel function (LOF) or aberrant gain-of-function (GOF) (*4–7*). LOF mutations cause type 1 long QT syndrome (LQTS) cardiac arrhythmia, a cause of sudden death. Less common GOF mutations are a cause of other arrhythmias such as short QT syndrome (SQTS) and atrial fibrillation (AF). We conducted a recent study of 50 different mutant forms of KCNQ1 in which the single mutation sites were all located in the voltage sensor domain (VSD) (*8*). We found that 31 of the mutants exhibited LOF, consistent with their association with LQTS. For a majority of these mutants, reduced cell surface channel levels were observed and were documented to result from the combined impact of proteasomal degradation and impaired trafficking to the cell membrane. For this majority of LOF mutants the most common underlying defect resulting in degradation or mis-trafficking was seen to be destabilization of the folded state of the channel. These results provided a well-documented example of what is very likely a general principle regarding disease-promoting LOF mutations in human membrane proteins—that the most common cause of LOF is mutation-induced destabilization of the native conformation (*9*).

The fact that LQTS-linked KCNQ1 LOF is commonly caused by reduced surface channel levels led us to hypothesize that mutations resulting in aberrant gain of KCNQ1 function might in some cases result from an increase in its cell surface expression, which could arise from increased overall expression, decreased channel degradation, and/or enhanced surface trafficking efficiency. In this paper, we survey known disease-linked GOF mutant forms of KCNQ1 and report that one of these mutants, R231C, is indeed a “super-trafficker”. We also report studies of this R231C mutant that illuminate the physicochemical basis for this super-trafficking trait and reveal how this mutant results in a complex phenotype under human disease conditions.

## RESULTS

### R231C KCNQ1 exhibits super-trafficking behavior

We first examined the expression of 15 previously identified mutations that cause a GOF in KCNQ1 (*10–29*), as summarized in table S1. GOF mutations were introduced into cDNA encoding an epitope-tagged form of KCNQ1 bearing a myc epitope in an extracellular loop connecting the S1 and S2 transmembrane segments, a modification that has no impact on channel folding, trafficking or function (*30*). Each GOF variant was then transiently expressed in HEK293 cells, and the expression of each variant was measured by FACS-based quantification of anti-myc immunostaining. Two distinct fluorescently-tagged anti-myc antibodies were used to differentially label cell surface and intracellular KCNQ1 levels (*8*). From these measurements it was possible to calculate the cell surface trafficking efficiency: (surface expression/total expression) X 100. Results are shown in Figure 1. Fig. 1A shows that GOF mutant forms of KCNQ1 exhibit a wide range of surface expression levels relative to WT, ranging from about 10% for G229D to a remarkable 500% for R231C. Because of its unusual “super-trafficking” properties we focused all our subsequent work on.investigating R231C. The 5-fold increase in surface expression of R231C reflects the combined effects of its 1.7-fold higher level of total expression relative to WT (Fig. 1B) and its 3-fold greater surface trafficking efficiency (Fig. 1C).

**Figure 1.**
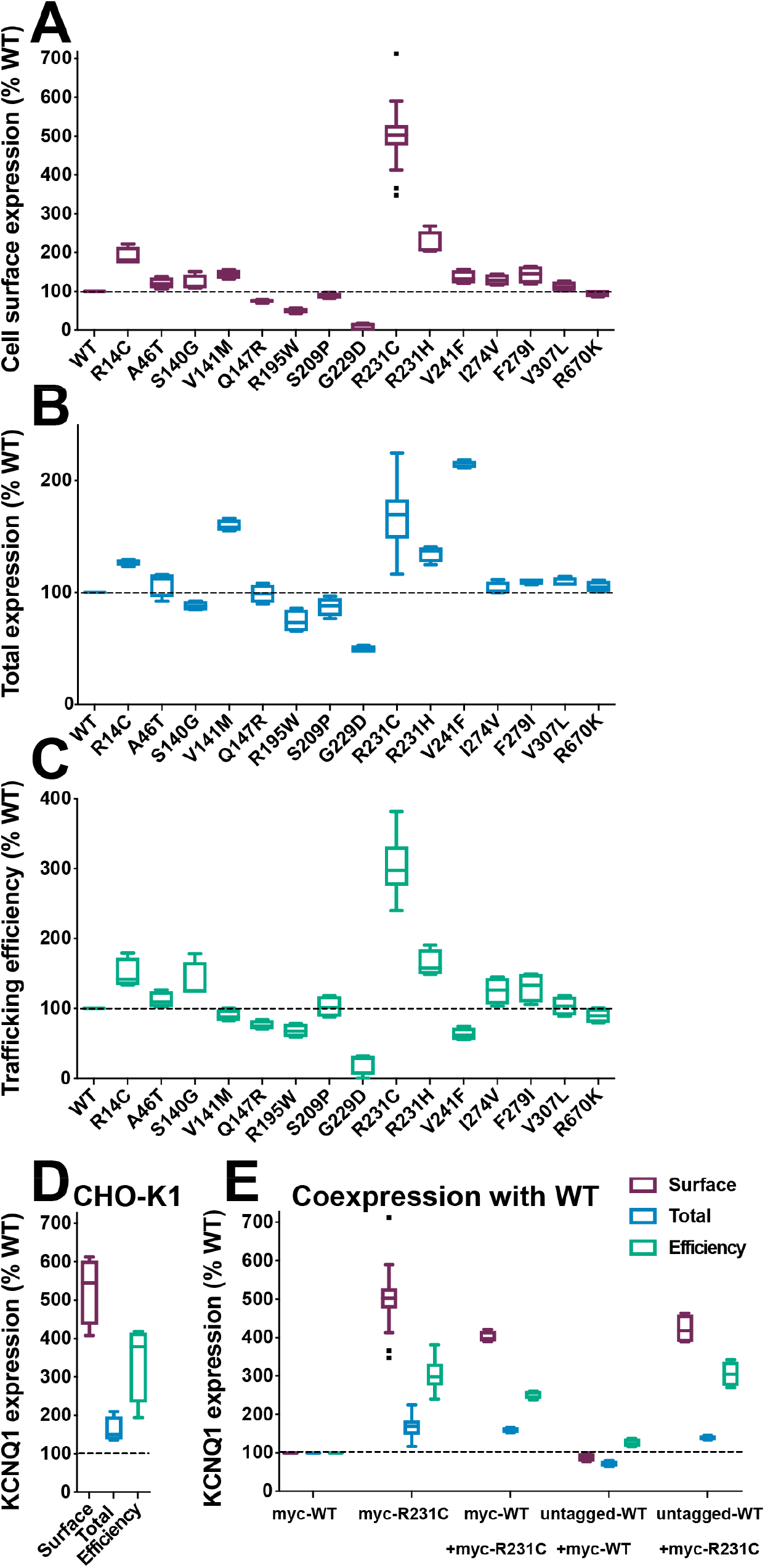
KCNQ1 GOF mutants display varied total and cell surface expression levels and trafficking efficiencies. The cell surface expression level (**A**), the total cell expression level (**B**), and surface trafficking efficiency (**C**) of GOF mutants of KCNQ1 were quantitated in transiently transfected HEK293 cells by flow cytometry. The WT trafficking efficiency is 15.4% (± 0.7%). (**D**) R231C also super-traffics in CHO-K1 cells. (**E**) HEK293 cells were transfected with 0.5 μg WT or R231C DNA (homozygous conditions) or were co-transfected with 0.25 μg WT and 0.25 μg R231C DNA (heterozygous conditions). Data are shown as percent of WT after correction for nonspecific staining of mock transfected cells. The results represent 22 experiments for R231C in HEK293 cells and 4 experiments for all other samples. 2500 single cell measurements were made per experiment. All panels present the data as Tukey box plots are presented where the central line denotes the median, the box denotes quartiles, and the whisker encompasses values within 1.5 times of the interquartile range.

For R231C, the trafficking results of Fig. 1 were not expression system-dependent; similar results were obtained when CHO-K1 cells (Fig. 1D) were used for channel expression instead of HEK293. We also examined whether co-expression of R231C affected WT channel surface trafficking. Transfection of myc-tagged 5-fold higher levels of KCNQ1 protein at the plasma membrane than following transfection of myc-tagged WT channel. When myc-R231C was co-transfected with myc-WT (same amount of total myc-R231C + myc-WT cDNA as myc-R231C cDNA in the previous experiments), the detected myc-tagged protein is 4-fold higher than WT channel expressed alone. (Fig. 1E). This indicates that co-assembled WT/R231 KCNQ1 heterotetramers also super-traffic.

Figures. 2A and 2B give the single cell trafficking efficiencies for WT and R231C expressing cells as a function of total cellular expression. The average trafficking efficiency of WT KCNQ1 is 15.4% ± 0.7% (S.E.M.).while the trafficking efficiency for R231C is much higher with an average trafficking efficiency of 42.8% ± 2.0%. When these data were binned into 10 equally sized cohorts (ranked by single cell total expression), the single cell trafficking efficiency for both WT and R231C KCNQ1 was seen to be inversely proportional to total single cell expression (Fig. 2C). However, while R231C trafficking efficiency does decline with higher expression levels, the drop is not as steep as for WT. Figs. 2E and 2F provide further clarification by showing that as the total expression of both WT and R231C increase, levels of both surface-trafficked and cell-internal KCNQ1 increase, but that for WT the internally trapped fraction grows especially more steeply. This suggests that R231C undergoes forward trafficking through the protein folding quality control system of the endoplasmic reticulum more readily than the WT channel.

**Figure 2.**
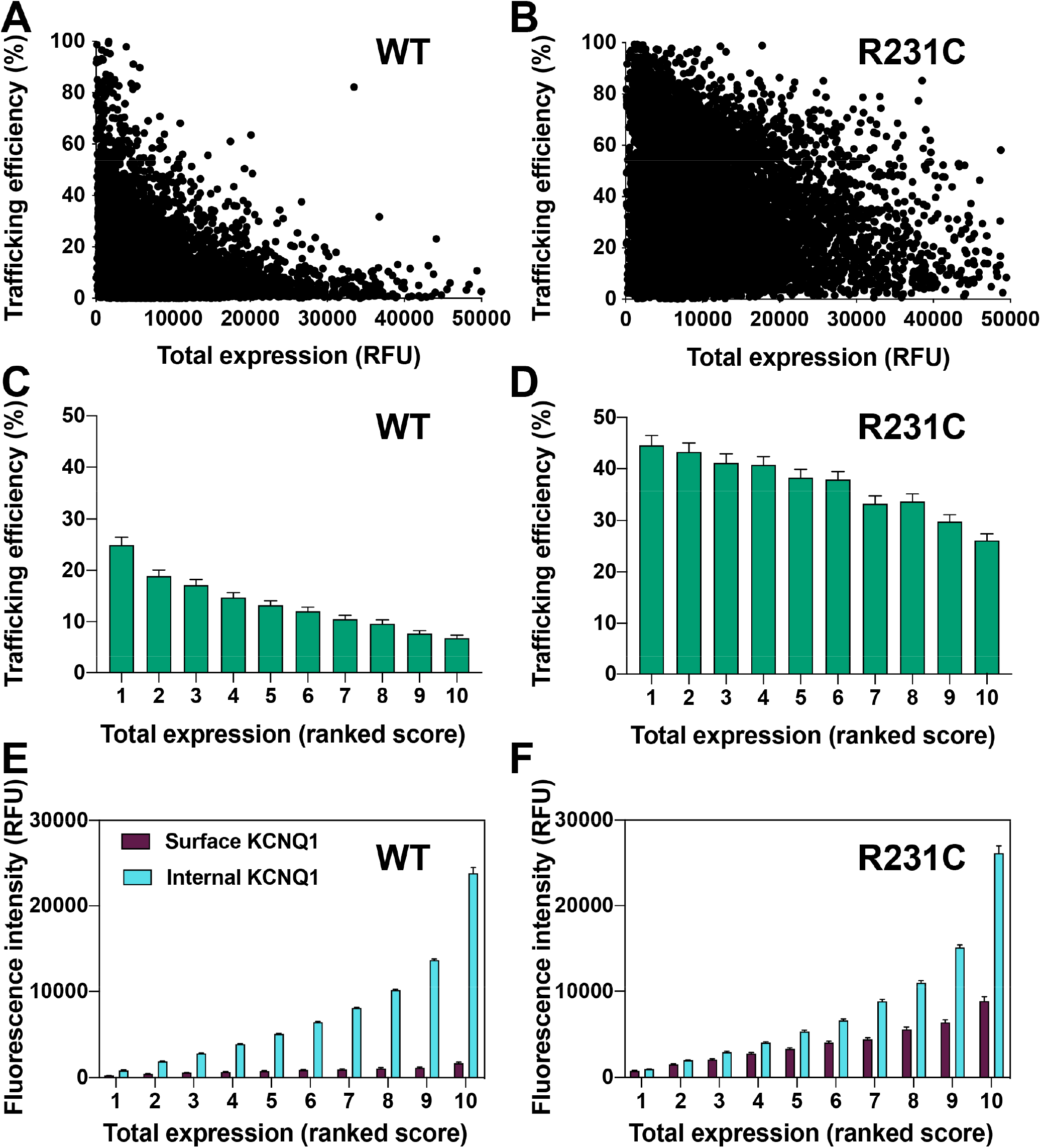
Single cell trafficking properties are analyzed for WT versus R231C KCNQ1 in HEK293 cells. The internal expression and trafficking efficiency of WT and R231C were determined for each of 7500 cells from three independent biological replicates. Cells that exhibited a KCNQ1 trafficking efficiency of zero were excluded from this analysis. (**A** and **B**) Plots of single cell KCNQ1 trafficking efficiency (where trafficking efficiency = (KCNQ1 surface expression level for each cell / KCNQ1 total expression for each cell) X 100. Results were ordered from lowest to highest KCNQ1 total expression and then binned into 10 groups of 750 each, designated 1 (for the 10% of cells exhibiting the lowest total KCNQ1 expression levels) to 10 (for the 10% with the highest total KCNQ1 expression levels). The KCNQ1 trafficking efficiency (**C** and **D**) and surface or internal expression levels (**E** and **F**) have been plotted as a function of the 10 bins ranked based on total expression levels. The bar height is the mean value and the error bar shows the 95% confidence interval for each group.

### Super-trafficking of R231C KCNQ1 is unrelated to disulfide bond formation

We considered the possibility that super-trafficking could be related to disulfide bond formation between the introduced Cys231 in S4 and the only native cysteine residue in its general vicinity, C136, located in the S1 transmembrane helix. As shown in fig. S1, the C136A/R231C double mutant exhibits enhanced cell surface expression similar to R231C, ruling out intramolecular disulfide bond formation as a causative or contributing factor to super-trafficking of this mutant. We also examined the consequences of mutating G229, I230, F232, or L233 to cysteine (instead of R231). None of these KCNQ1 mutants exhibited super-trafficking (fig. S1) indicating that the super-trafficking effect of mutation to cysteine is position-specific.

### Mutations of R231 to other amino acids lead to a range of KCNQ1 trafficking efficiencies

We next quantitated the expression and trafficking of a series of thirteen R231 mutants (Figure 3). R231C exhibits the highest level of expression, as well as the highest surface expression and trafficking efficiency. However, other mutants exhibit super-trafficking to a lesser degree, with trafficking efficiencies on the order of 2 to 3-fold higher than WT being observed for R231M, R231V, R231L and R231Y. Conversely, substitution of R231 to more polar residues (lysine and glutamate) results in even lower total expression and surface expression levels than WT.

**Figure 3.**
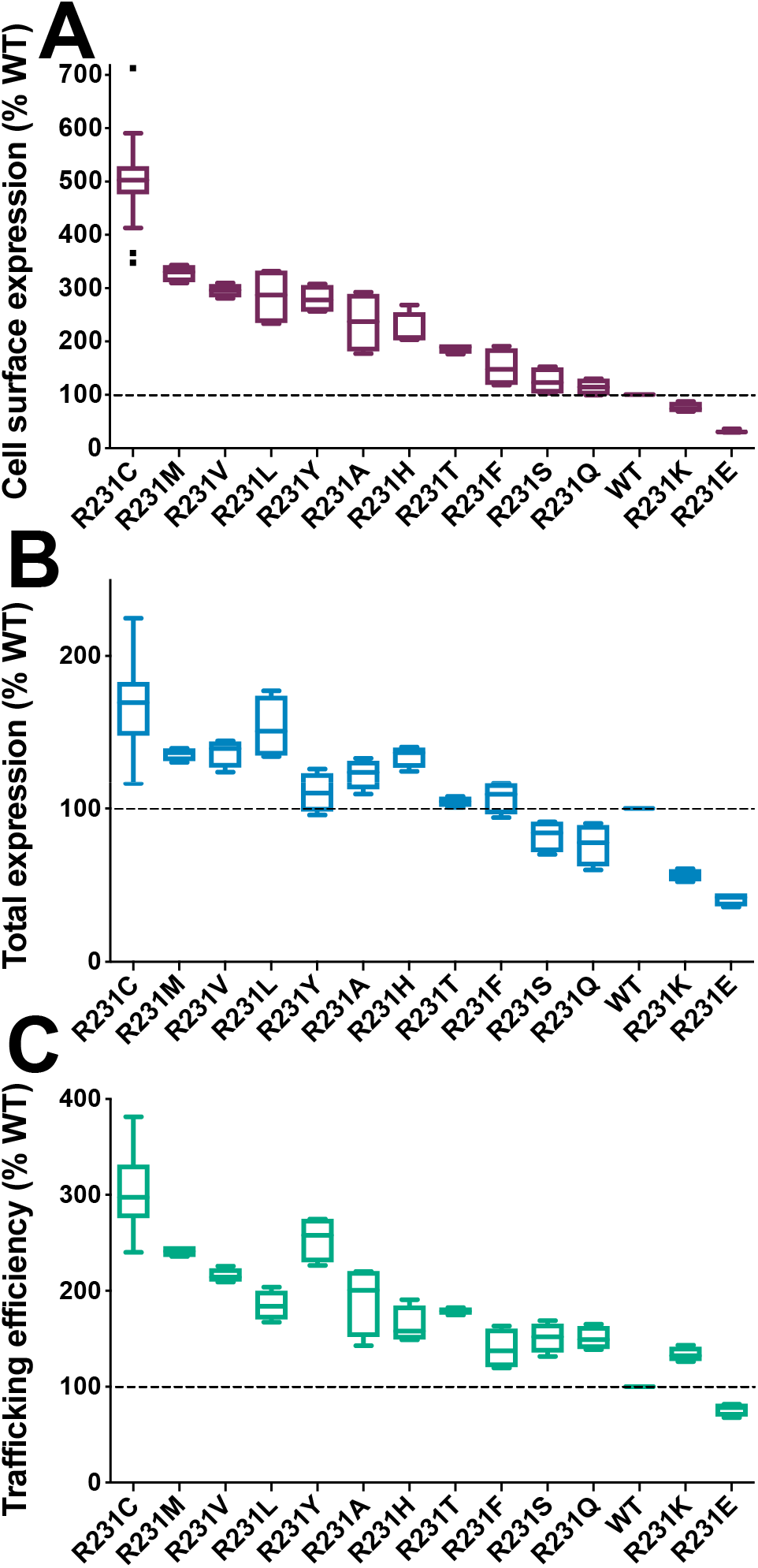
Mutations of R231 to other amino acids lead to a range of KCNQ1 trafficking efficiencies in HEK293 cells. See the legend of Figure 1 for additional details.

### Super-trafficking of R231 mutants is related to the membrane integration of KCNQ1 S4 helix

R231 is located within the S4 helix, a key element of the voltage sensor domain of KCNQ1 and other voltage-gated ion channels. The S4 helix contains positively charged residues (Arg or Lys), which are uncharacteristic for a transmembrane helix (*31–33*). While these Arg and Lys residues are critical to the voltage-sensing function of S4, they also compromise the co-translational membrane integration of this helix (*31, 34–37*). We hypothesized that the increased hydrophobicity for substitutions at R231 might enhance membrane integration of S4, which in turn increases the folding efficiency of the channel. We first examined this issue by comparing the trafficking results of Fig. 3 to the projected effects of these mutations on the membrane integration of S4 according to the Hessa/White/Von Heijne “biological hydrophobicity scale”, which is a knowledge-based energy scale for the translocon-mediated membrane integration of nascent transmembrane helices (*38, 39*). Figure 4A-4C shows that the surface trafficking, total expression, and trafficking efficiency of R231 variants are highly correlated with the projected effects of these mutations on the transfer free energy of the S4 helix. Overall, the trafficking of the 13 mutants examined in this work appear closely related to their impact on the hydrophobicity of S4. However, as seen in Fig. 4A-4C, R231C is an outlier that expresses and traffics even higher than expected. Indeed, its surface expression level is roughly two-fold higher than expected based on the best linear fit of the stie 231 mutant series expression levels to the Hessa/White/von Hiejne ΔΔG values.

**Figure 4.**
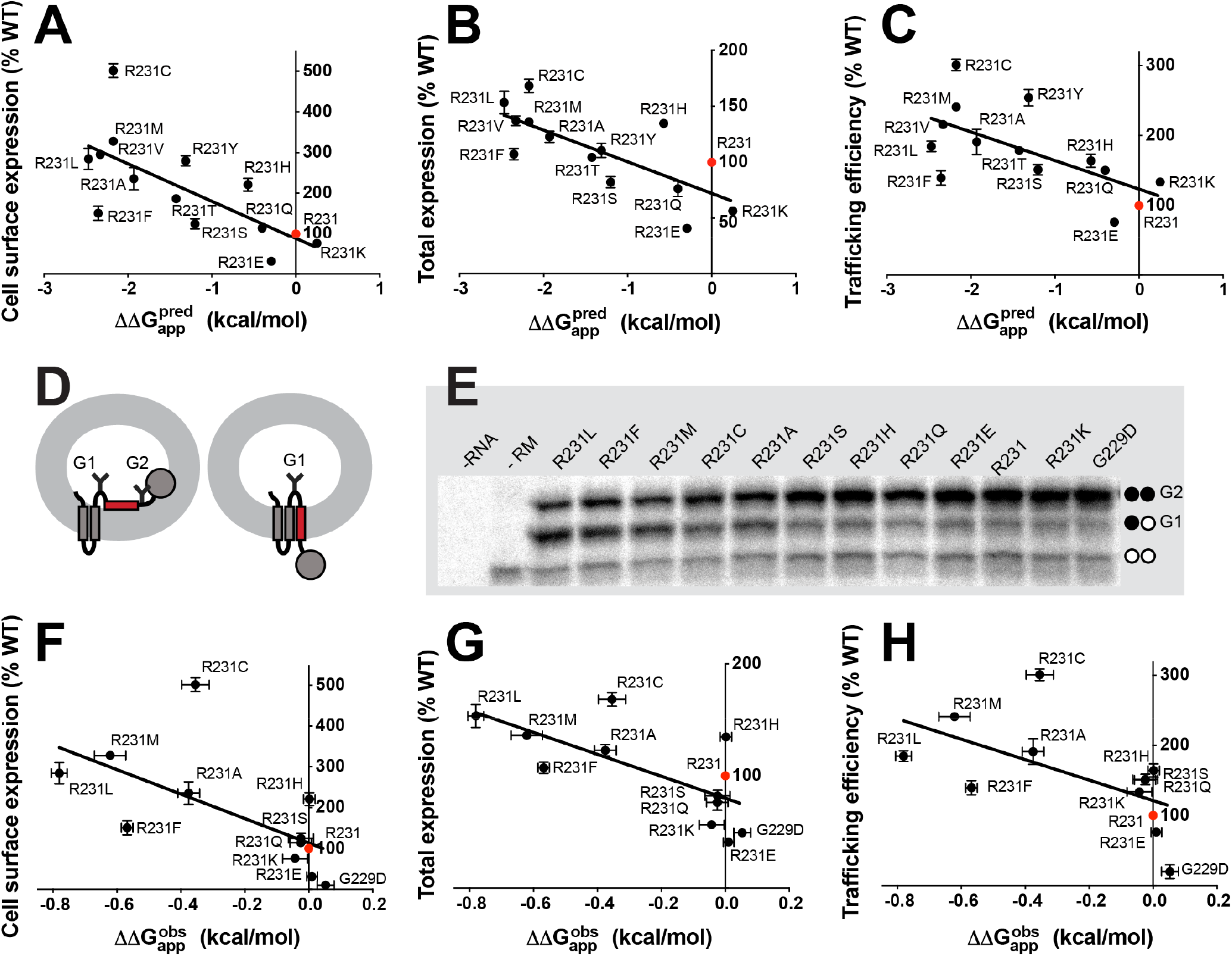
The super-trafficking of R231C is related to increased membrane integration of the KCNQ1 S4 helix. **(A**-**C)** The Hessa/White/Von Heijne biological hydrophobicity scale helps to explain the trafficking of some 13 different KCNQ1 site 231 mutants in HEK293 cells. Here, 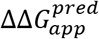 is the predicted (http://dgpred.cbr.su.se/) apparent free energy difference between correct integration of the site 231-containing KCNQ1 S4 segment into ER membrane by the Sec61 translocon versus the fraction that is predicted not to integrate into the membrane. (**D**-**H**) *In vitro* translation results for site 231 mutants of the isolated KCNQ1 S4 segment show a generally excellent correlation with the expression of the same mutant series in full length KCNQ1 expressed in HEK293 cells. The assay used a Lep construct by replacing its H-segment with the KCNQ1 S4 segment (red). Chimeric Lep constructs encoding KCNQ1 mutants were transcribed and translated *in vitro* with canine rough microsomes (RM) and analyzed by SDS-PAGE. Doubly (G2) and singly (G1) glycosylated Lep proteins were quantified. Unglycosylated products are labeled with two white dots. Control reactions were performed without adding RNAs or without adding RMs. 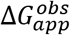 values were determined from at least three independent biological replicates and compared to that for WT to generate the reported 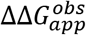. Data plotted is the mean value and the error bar is S.E.M..

Based on these observations, it is unclear whether the deviation in expression caused by the R231C mutation stems from its effects on the membrane integration of S4, its impact on the structural properties of the full-length channel, or some combination of the two. To examine R231C effects on S4, we measured the membrane integration of S4 variants in the context of a chimeric leader peptidase protein, as previously described (*38*). Lep proteins were produced by *in vitro* translation in microsomes, and the membrane integration efficiency of each S4 variant was inferred from the relative abundance of the singly (S4 ends up in the membrane) and doubly (S4 ends up in the lumen) glycosylated forms of the protein (Fig. 4D). Consistent with observations for other S4 segments (*35, 37*), the S4 segment of KCNQ1 exhibits a marginal propensity to partition into the membrane (ΔG_app_ = 0.56 ± 0.03 kcal/ mol Fig. 4E). The measured change in the apparent free energy of membrane integration (ΔΔG_app_) caused by these mutations is small relative to predicted values, which likely reflects the limited dynamic range of this assay (*38, 40*). Despite this limitation, the introduction of hydrophobic side chains at R231 clearly increases the membrane integration efficiency of S4 (Fig. 4E). Furthermore, measured ΔG_app_ values are inversely correlated with the surface expression, total expression, and trafficking efficiency of full length KCNQ1 mutants (Fig. 4F-4H). Importantly, the experimental R231C ΔG_app_ is again seen to be an outlier relative to the best linear fits of the trafficking data (Fig. 4F-4H), indicating that its elevated expression level and cell surface trafficking efficiency can be only partly accounted for by its more efficient membrane integration by the translocon.

### Energetic coupling of the R231C and F166 side chains is a second factor that contributes to the super-trafficking behavior of R231C KCNQ1

We sought to determine if interactions between the R231C side chain and those of proximal residues in the folded conformation of the KCNQ1 voltage sensor domain contribute to channel super-trafficking. To identify the structural interactions that influence KCNQ1 trafficking, we quantitated the surface expression of a series of single and double mutants that selectively perturb possible contacts of interest. Our strategy was based on the notion that surface expression should be attenuated by single mutations that disrupt a critical interaction, yet restored by the introduction of the reciprocal (swapped sites) mutation. To avoid complications from possible disulfide bond formation, we sometimes used R231M as the parent mutant, with key results then being validated by repeating measurements using R231C.

We selected potential interacting sites with R231C using the available experimental structures for the fully activated KCNQ1 channel (*41*) and intermediate-activated voltage sensor domain (*42*), as well using a recently developed inactive state model (*43*). Based on these structures, C231 is predicted to be to close or in contact to sites 143,144, 212, 278, 281, and 299 in the fully activated state, to sites 140, 163, and 209 in the intermediate state, and to sites 129, 133, 166, and 167 in the inactive state (fig. S2). Figure 5A shows that reciprocal mutations of the side chain of R231C or R231M with the WT side chains of the following residues led to a complete or near-complete loss of super-trafficking: S140, S143, T144, F167, S209, Y278, Y281, and Y299. While it is possible that these side chains may interact with residue 231 in certain conformational states of the channel, these results suggest any such interaction does not contribute to the super-trafficking trait of the R231C channel. This inference was definitively confirmed by additional mutagenesis measurements involving for these sites as shown in fig. S3.

**Figure 5.**
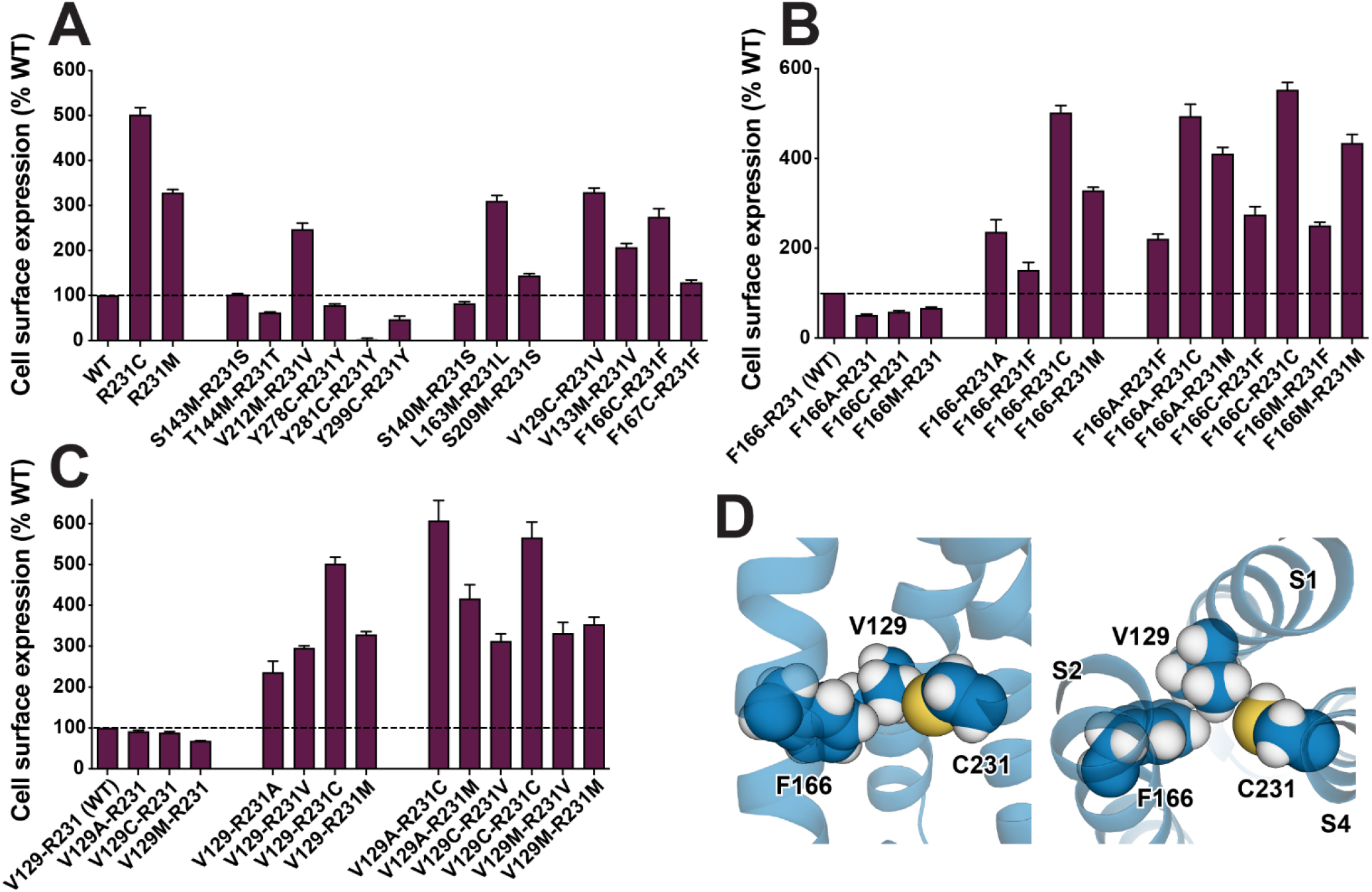
Double mutant analysis suggests that residues V129, F166, and C231 are energetically coupled in a way that promote super-trafficking of R231C KCNQ1. (**A**) Double reciprocal (swapped-site) mutagenesis for site 231 in parent mutants R231M or R231C and various potential interacting sites. The total expression level and cell surface trafficking efficiency of swapped double mutants were measured. (**B** and **C**) Follow-up mutagenesis experiments to the data in panel A for sites 166 and 129, respectively. Data are shown as percent of WT after correction for nonspecific staining of mock transfected cells. Bar height indicates mean value, where the error bar is S.E.M. of at least three independent biological replicates. (**D**) Tri-partite interaction of the energetically-coupled side chains of C231, V129, and F166 in the inactive state structure (*43*) of the KCNQ1 VSD. It is seen that while C231 does not directly contact F166, V129 forms an intimate bridge between C231 and F166. Both side view (left) and top view (right) are shown.

Fig. 5A also shows several possible R231C-interactor sites that retain considerable super-trafficking capability when mutationally swapped with site 231 Cys or Met: V129, V133, L163, F166, and V212. Follow-up mutagenesis studies were conducted to explore the possibility of trafficking-impacting energetic linkage between the side chains of these sites and that of site 231 (Fig. 5B, 5C and fig. S3). For sites V133, L163 and V212, the additional mutagenesis data (fig. S3) does not reveal significant energetic linkage: non-conservative mutation of each of these sites in the presence of R231C or R231M either resulted in no impact or in a negative impact on trafficking that was in no case rescued by a subsequent swap of that residue with the Cys231 or Met231. Instead, the impact of mutations at these sites on trafficking is independent and additive with the impact of the mutations at site 231. However, for F166 and V129 the data of Figs. 5B and 5C is consistent with a trafficking-impacting energetic linkage between its aromatic side chain and either the R231C or R231M side chains. Fig. 5B shows that mutation of F166 to Ala, Cys, or Met in WT (R231) KCNQ1 results in reduced trafficking. However, when mutation of F166 to any of these amino acids is combined with the mutation of R231 to Ala, Cys, Met, or Phe the result is super-trafficking for each double mutant (Figs. 5A and 5B). Sites R231C and F166 are evidently energetically coupled in a manner that helps increase trafficking efficiency. A similar pattern is seen for V129, indicating that this site is also energetically coupled to R231C (Fig. 5C). From the available 3D structures of the KCNQ1 VSD we see that the side chain of R231C is close to the side chains of V129 and F166 only when the VSD populates the inactive state (Fig. 5D and fig. S2). When R231 is computationally mutated to cysteine, its side chain interacts directly with V129, while the side chain of V129 is in intimate direct contact with F166. Coupling between the R231C site and V129 is, therefore, direct, while coupling between R231C and F166 is mediated by V129.

### Functional analysis validates energetic coupling between residues 231 and 166 and also shows that R231C and related mutants are both constitutively active but have reduced activity relative to the WT channel

Both WT and selected super-trafficking KCNQ1 mutants were functionally analyzed using a CHO-K1 cell line stably-expressing KCNE1. KCNE1 is a channel accessory subunit that pairs with KCNQ1 to form the channel complex responsible for the cardiac I_Ks_ current (*44*). Figure 6A shows that the super-trafficking KCNQ1 mutants R231C, R231M, and F166M/R231F exhibited constitutive channel activity in the −80 mV to +60 mV voltage-range. The same observation is made for R231F, which traffics more efficiently than WT but by only ca. 50%, indicating that for R231 mutations constitutive activity and super-trafficking are not correlated. The results for these R231 mutants are in contrast to those for WT and F166M forms of KCNQ1 (Fig. 6A), which are voltage regulated—opening only upon membrane depolarization, with a V_1/2_ of roughly +30 mV. Fig. 6B depicts the average current-voltage (IV) relationships measured after 2s pulses from CHO_KCNE1 cells transiently transfected with KCNQ1 WT, F166M or R231 mutants. The whole cell current density measured at +60 mV for R231C is similar to that for WT, whereas R231F exhibits significantly larger current. In contrast, R231M, F166M and F166M-R231F exhibit smaller current density than the WT channel at +60 mV. Fig. 6C shows the current-density measured for each mutant KCNQ1 channel (Fig. 6B) normalized to the cell surface expression levels relative to WT (see Fig. 5). By analyzing the current measured at +60 mV, this plot reveals that R231C and R231M exhibit decreased function relative to WT. While these mutants are constitutively open, their activity per channel protein at the membrane is only on the order of 20% that of WT. In contrast, the R231F and F166M mutants have similar activity at +60 mV when compared to the WT channel, while the F166M/R231F double mutant is only 20% of WT. This result indicates that energetic coupling between certain pairs of side chains for residues 166 and 231 can contribute not only to super-trafficking but to a dramatic reduction in channel function by either decreasing single-channel conductance, lowering open state probability, or both. We again note that sites 166 and 231 are in proximity (bridged by contact with V129, Fig. 5D) only when the channel populates the inactive state, suggesting that the decrease in channel activity seen for R231C and R231M is due to reduced open state probability. These mutations likely stabilize the non-conductive inactive state, suggesting an equilibrium between states in which the inactive-to-active state population ratio is roughly 4:1 in favor of the inactive state.

**Figure 6.**
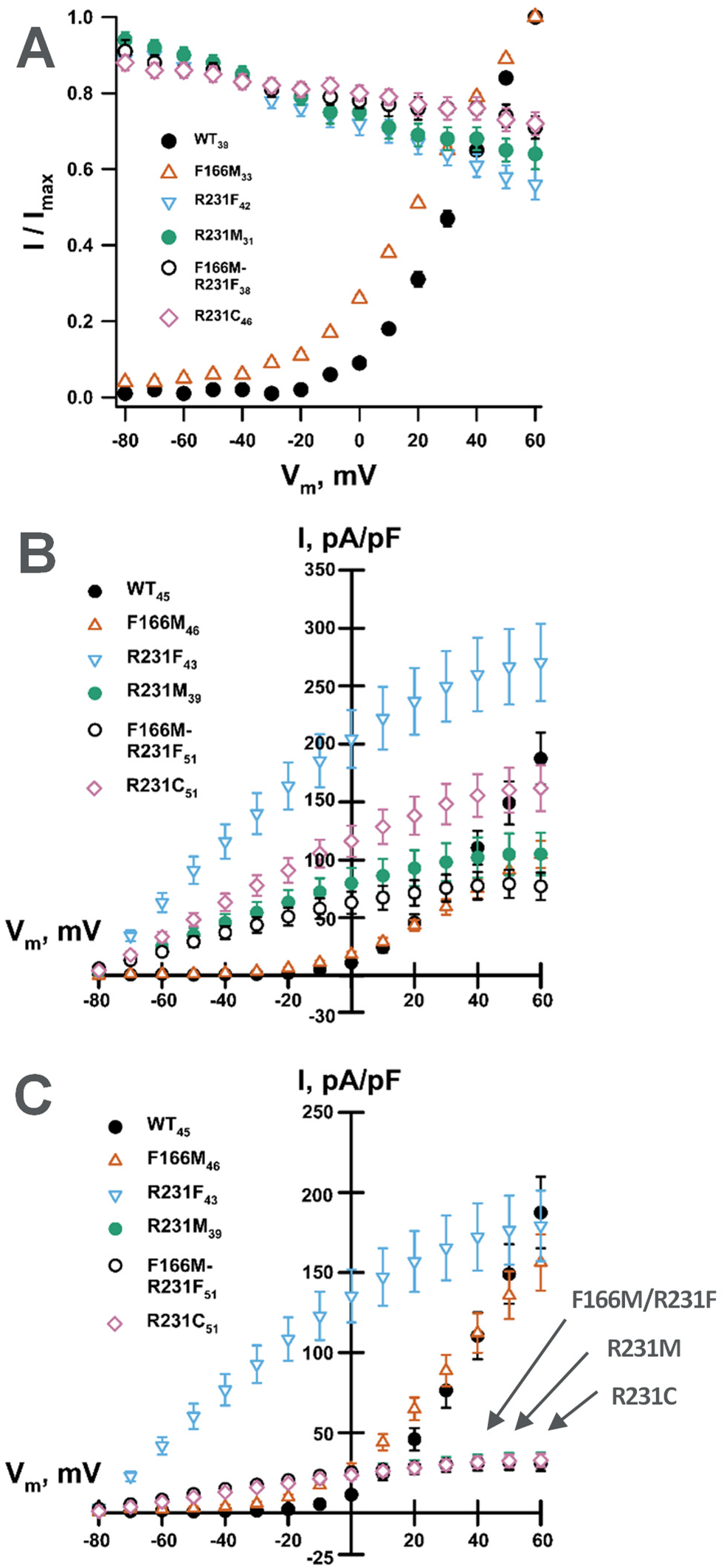
Electrophysiological functional analysis validates energetic coupling between residues 166 and 231 and shows that R231C and related mutants are constitutively active but also have reduced activity relative to the WT channel. CHO-K1 cells stably expressing KCNE1 were electroporated with plasmids containing either WT or mutant KCNQ1 cDNA. (**A**) Voltage-dependence of the activation curves obtained from whole-cell currents recorded from CHO_KCNE1 cells transiently expressing WT or mutant KCNQ1 channels. (**B**) Current density-voltage relationships measured in CHO_KCNE1 cells transiently expressing WT or mutant KCNQ1 channels. (**C**) Current density-voltage relationships measured in CHO_KCNE1 cells transiently expressing WT or mutant KCNQ1 channels and normalized based on the measured quantity of channel protein at the plasma membrane. Whole-cell currents were recorded using automated patch clamp, normalized by membrane capacitance and the number of recorded cells (n) is labeled next to each mutant.

Overall, the results of Fig. 6 are consistent with previous observations (*10, 17, 45*) that mutations of R231 to neutral residues result in constitutive (voltage-independent) channel activation. However these results also provide new insight that a number of these very same mutations, including R231C, exhibit dramatically reduced channel activity relative to WT. Moreover, the observation that mutation of site 231 to cysteine leads to energy coupling of this site to sites V129 and F166 to enhance channel trafficking is consistent with the proximity of the side chains for all three of these residues, which occurs only in the inactive state. This leads to a complete model for how R231C results in disease-associated KCNQ1 dysfunction, as described below.

## DISCUSSION

### Defects in the R231C KCNQ1 channel are three-fold

Dominant missense mutations in the human *KCNQ1* gene sometimes result in the formation of defective channel proteins. While a majority of the known KCNQ1 disease variants exhibit channel LOF that results in long QT syndrome arrhythmia, a smaller set of variants result in what is thought to be aberrant GOF, resulting in short QT syndrome or familial atrial fibrillation (*4–7*) (table S1). The R231C mutant has previously been included in this classification because it is known to be constitutively open at all transmembrane potentials (*10, 17, 45, 46*) This is in contrast to healthy conditions for the cardiac action potential, where gating of the KCNQ1 channel in complex with its modulatory submit KCNE1 is voltage-regulated. The results of this work revealed two additional traits that set it apart from WT and other GOF mutant forms. One is that while it is constitutively active, the activity per channel protein of R231C in complex with KCNE1 is only 20% that of the WT KCNQ1/KCNE1 complex. The other is that this mutant is much more abundant at the plasma membrane than the WT channel (Fig. 1), an observation that holds even under R231C/WT heterozygous conditions (Fig. 1E). This increase in channel levels at the plasma membrane offsets the 5-fold decrease in channel activity. Accordingly, arrhythmia caused by the R231C KCNQ1 can be regarded as being due both to super-trafficking of the channel that partially cancels out reduction in total cell KCNQ1 current resulting from reduced channel activity, plus channel dysregulation in the form of voltage-independent constitutive conductance. Our results provide clarity as to how R231C KCNQ1 results in pleiotropic arrhythmic phenotypes including AF, LQTS, and fetal bradycardia (*10, 17, 46, 47*). Depending on exactly in which part of the heart heterozygous expression of KCNQ1 occurs, the balance between super-trafficking and reduced activity condition very likely varies, such that R231C-like I_Ks_ may locally exhibit either net LOF properties (promoting LQTS) or dysregulated GOF behavior (promoting superimposed AF).

### Structural basis for R231C super-trafficking

We observed that the mutation of R231 to hydrophobic residues generally increases the abundance of KCNQ1 at the cell surface, with the results for R231C being especially striking. The “super-trafficking” activity of this mutant reflects both the ~1.7-fold higher total expression level of R231C KCNQ1 and its 3-fold higher surface trafficking efficiency relative to WT. Our results suggest this trait arises from two contributing factors. First, about half of the super-trafficking can be attributed to enhancement by this mutation of the efficiency of translocon-mediated membrane integration of the S4 helix. The WT R231 residue contributes to the polarity of the S4 transmembrane segment and its resulting inefficient membrane integration. Despite this inefficiency, this residue is highly conserved due to its indispensable role in transmembrane voltage sensing. Though the introduction of hydrophobic amino acids at this position alters channel activation, we show that this also increases the translocon mediated-membrane integration of S4 in a manner that correlates with the increased expression and trafficking efficiency of the full-length channel (Fig. 4). These findings parallel recent investigations of the co-translational folding of rhodopsin, where it was found that its expression and trafficking were highly sensitive to mutations that alter the hydrophobicity of a polar transmembrane domain containing functionally constrained polar residues (*48, 49*). However, the R231 mutations characterized herein demonstrate that the inefficiency of translocon-mediated membrane integration can cut both ways. Though mutations that introduce polar residues within transmembrane domains can cause a pathogenic LOF by compromising folding(*50, 51*), our findings also suggest hydrophobic mutations that enhance folding and expression can also potentially result in a toxic GOF. Given that action potentials hinge on a precise sequence of controlled ionic fluxes, native expression levels very likely have been tuned by evolution to factor in inefficiency that arises from frustrated regions that are critical for function.

In addition to the effect of the R231C mutation on the efficiency of co-translational folding, we find that this mutation also enhances an energetic coupling in the inactive state that enhance the expression of the full-length channel. To identify the structural basis of this enhanced expression, we explored the proteostatic effects of mutations that perturb adjacent residues in combination with R231C. It has previously been shown for KCNQ1 and other membrane proteins that both total expression levels and surface trafficking is dependent on the stability of the folded form of protein, with more stable forms trafficking to the cell surface with increased efficiency (*8, 9, 52*). The mutagenesis studies of this work confirmed that introduction of a cysteine at site 231 leads to energetic coupling with V129 and F166 that shifts the voltage sensor equilibrium to favor the inactive state relative to the activated state by 4:1, leading to the reduced conductance seen for this mutant. This C231/V129/F166 tri-partite interaction evidently also results in the extra 2-fold increase in surface trafficking beyond what can be expected based solely on the relative hydrophobicity of cysteine relative to arginine.

## CONCLUSIONS

This work documents that some disease mutations of membrane proteins contribute to human disease by promoting dramatically enhanced trafficking of the mutant protein to its correct destination membrane in the cell. This class of mutation is rare among the roughly 50 disease mutant forms of KCNQ1 for which quantitative trafficking data is now available. Documentation of this super-trafficking trait in this work establishes a “too much of a good thing” paradigm that seems likely to be seen in disease mutant forms of other membrane proteins.

## MATERIALS AND METHODS

### Plasmids and mutagenesis

The untagged and c-myc tagged human KCNQ1 DNA (GenBank accession number AF000571) were engineered in the mammalian expression vector pIRES2-EGFP as previously described (*8*). The c-myc tag (EQKLISEEDL) was inserted into the extracellular S1-S2 linker between residues Glu146 and Gln147. Human KCNE1 (L28168) was subcloned into the pcDNA3.1(+) vector for biochemical experiments and into pcDNA5/FRT to generate the CHO_KCNE1 cell lien used for functional analysis of KCNQ1 mutants as previously described (*8, 53*). InFusion HD Cloning (Takara Bio, Mountain View, CA) was used to introduce the sequence of the WT S4 domain of KCNQ1 in place of the H-segment of a modified leader peptidase (Lep) gene, as previously described (*39*). Mutations in KCNQ1 were introduced by QuikChange site-directed mutagenesis using untagged or c-myc tagged KCNQ1 plasmid as the template. Mutations in the S4 segment of Lep were generated using the WT S4 Lep as the template. The coding regions of all constructs were verified by Sanger sequencing.

### Cell culture

Human embryonic kidney cells (HEK293, CRL-1573) and Chinese hamster ovary cells (CHO-K1, CRL-9618) were purchased from American Type Culture Collection (Manassas, VA). HEK293 cells were grown in Dulbecco’s modified Eagle’s medium (DMEM) supplemented with 10% fetal bovine serum (FBS), 10 mM HEPES, 100 units/ml penicillin, and 100 μg/ml streptomycin at 37 °C in 5% CO_2_. CHO-K1 cells were cultured in F-12 nutrient mixture medium supplemented with 10% FBS, 50 units/ml penicillin, 50 μg/ml streptomycin, and 2 mM L-glutamine at 37 °C in 5% CO_2_. As previously described (*45*), CHO-K1 cells stably expressing KCNE1 (CHO_KCNE1) were made using KCNE1 in pcDNA5/FRT vector and generated using the FLP-in™ system (Thermo Fisher Scientific, Waltham, MA) following the manufacturer’s instruction. CHO_KCNE1 cells were maintained under dual selection with Zeocin (100 μg/ml) and hygromycin B selection (600 μg/ml).

### Flow cytometry

Unless otherwise stated, HEK293 cells were used for the flow cytometry study of KCNQ1 total and surface expression levels. Cells were plated into six-well plates and the next day transiently transfected with 0.5 μg WT or mutant myc-KCNQ1, each well at 30-50% confluence. Transfection was performed using Fugene 6 transfection reagent (Promega, Madison, WI). When WT and mutant myc-KCNQ1 were co-transfected, the ratio of DNA used was 1:1 (0.25 μg: 0.25 μg). When KCNE1 in pcDNA3.1(+) vector and myc-KCNQ1 were co-transfected, the ratio of DNA used was also 1:1 (0.5 μg: 0.5 μg).

Approximately 48 h after transfection, cells were prepared for flow cytometry quantitation. As previously described (*8, 52*), cells were prepared using the Fix & Perm cell fixation and cell permeabilization kit (Thermo Fisher Scientific) following the supplier’s protocol. All procedures were performed at room temperature. Briefly, cells were washed once with PBS containing 25 mM HEPES and 0.1% NaN_3_ (PBS-FC, pH 7.4), detached in PBS-FC buffer containing 0.5 mM EDTA and 0.5% BSA, and centrifuged at 300 × g for 5 min. Cells were then resuspended in 100 μl PBS-FC containing 5% FBS with 1:100 diluted PE-conjugated anti-myc (9B11) mouse monoclonal antibody (Cell Signaling Technology, Danvers, MA) and incubated for 30 min in the dark. 100 μl fixation medium was then added to each sample followed by incubation for 15 min to fix cells. Cells were washed once with PBS-FC containing 5% FBS. Cells were then resuspended in 100 μl permeabilization medium with 1:100 diluted Alexa Fluor 647-conjugated anti-myc (9B11) mouse monoclonal antibody (Cell Signaling Technology) and incubated for 30 min in the dark. Cells were washed once and fluorescence intensities were quantitated using a 4-laser Fortessa flow cytometer (BD Bioscience, San Jose, CA). To correct the fluorescence intensity difference of the two antibodies, cells expressing WT myc-KCNQ1 were fixed, permeabilized, and immunostained with either PE or Alexa Fluor 647-conjugated anti-myc antibody and the intensity ratio of the two antibodies was calculated.

Gating on GFP-positive cells was applied to select cells successfully transfected with the KCNQ1 plasmid. Fluorescence of mock transfected cells was used for background correction. Unless otherwise stated, the expression and trafficking efficiency of mutants were shown as percent of WT, where trafficking efficiency was defined as [(surface)_mutant_/(total)_mutant_]/[(surface)_WT_/(total)_WT_] X 100. The mean and S.E.M. values were calculated from at least three independent biological replicates. Data were analyzed and plotted using GraphPad Prism 8.2 software. Tukey box plot was used to illustrate data distribution and Student’s t-test was used to compare the differences between WT and mutants.

### In vitro translation of leader peptidase variants

A glycosylation based biochemical assay was used to measure the co-translational membrane integration efficiency of S4 variants. The WT and a series of Lep variants bearing mutations of interest within the KCNQ1 S4 segment were generated as described above. mRNA was synthesized from these plasmids using a RiboMAX SP6 RNA production kit (Promega). *In vitro* translation of the mRNA was carried out using rabbit reticulocyte lysate (Promega) supplemented with canine rough microsomes (tRNA probes, College Station, TX) and EasyTag ^35^ S-labeled methionine (PerkinElmer, Waltham, MA) for one hour at 30 °C. The products were separated on a 12% SDS-PAGE gel. The gel was dried for one hour at 60 °C, then exposed overnight on a phosphor imaging plate (GE Healthcare, New York, NY) and imaged on a Typhoon Imager (GE Healthcare). Doubly (*G2*) and singly (*G1*) glycosylated Lep proteins were quantified by densitometry in ImageJ software. Apparent transfer free energy values were calculated according to the following equation:

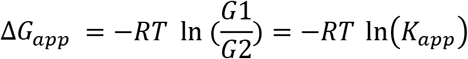

where Δ*G_app_* is the apparent transfer free energy for the transfer of the S4 helix from the translocon to the membrane, *R* is the universal gas constant, *T* is temperature, *G2* is the amount of doubly glycosylated protein, *G1* is the amount of singly glycosylated protein, and *K_app_* is the apparent equilibrium constant, as previously described (*38*). Δ*G_app_* are the average of at least three independent biological replicates and were compared to the Δ*G_app_* for WT to generate ΔΔ*G_app_* values.

### Electrophysiology experiments

Wild type and mutant untagged KCNQ1 were transiently transfected into CHO_KCNE1 cells using the Maxcyte STX system as previously described (*45*). Briefly, cells grown to 70-80% confluency were harvested using 5% trypsin. Cells were collected by gentle centrifugation (160 × g, 4 minutes), followed by washing the cell pellet with 5 ml electroporation buffer (EBR100, MaxCyte Inc.) and re-suspension in electroporation buffer at a density of 108 viable cells/ml. For each electroporation, plasmid encoding KCNQ1 variant (10 μg) was added to 100 μl cell suspension. The DNA-cell suspension mix was then transferred to an OC-100 processing assembly (MaxCyte Inc.) and electroporated using the CHO-PE preset protocol. Electroporated cells were grown for 48 hours at 37 °C in 5% CO_2_. Following incubation, cells were harvested, counted, transfection efficiency determined by flow cytometry and then frozen in 1 ml aliquots at 1.5×10^6^ viable cells/ml in liquid N_2_ until used in experiments.

Electroporated cells were thawed the day before an experiment, plated and incubated for 10 hours at 37 °C in 5% CO_2_. The cells were then transferred to 28 °C in 5% CO_2_ and grown overnight. Prior to the experiment, cells were passaged using 5% trypsin in cell culture media. Cell aliquots (500 μl) were used to determine cell number and viability by automated cell counting. Cells were then diluted to 200,000 cells/ml with external solution (see below), and allowed to recover 40 minutes at 15 °C while shaking on a rotating platform at 200 rpm.

Currents were recorded at room temperature in the whole-cell configuration by automated planar patch clamp using a SyncroPatch 768 PE (Nanion Technologies, Munich, Germany). Single-hole, 384-well recording chips with medium resistance were used in this study. The external solution contained (in mM) the following: NaCl 140, KCl 4, CaCl_2_ 2, MgCl_2_ 1, HEPES 10, glucose 5, with the final pH adjusted to 7.4 with NaOH. The internal solution contained (in mM) the following: KF 60, KCl 50, NaCl 10, HEPES 10, EGTA 10, ATP-K2 2, with the final pH adjusted to 7.2 with KOH. Whole-cell currents were not leak-subtracted. The contribution of background currents was determined by recording before and after addition of the IKs blocker JNJ-303 (4 μM). Only JNJ-303-sensitive currents were used for analysis.

Pulse generation and data collection were carried out with PatchController384 V.1.3.0 and DataController384 V1.2.1 software (Nanion Technologies). Whole-cell currents were filtered at 3 kHz and acquired at 10 kHz. The access resistance and apparent membrane capacitance were estimated using built-in protocols. Whole-cell currents were recorded from −80 to +60 mV (in 10 mV steps) 1990 ms after the start of the voltage pulse from a holding potential of −80 mV. The voltage-dependence of activation was calculated by fitting the normalized G-V curves with a Boltzmann function (tail currents measured at −30 mV). The number of cells (n) is given on the figure legends.

## SUPPLEMENTARY MATERIALS

**figure S1.** Super-trafficking of R231C KCNQ1 is unrelated to disulfide bond formation.

**figure S2.** Location of residues used for double mutant analysis of interactions involving C231 is illustrated in KCNQ1 structures.

**figure S3.** Double mutant analysis has been performed for sites that were deemed not to be energetically coupled to the side chain for site 231.

**table S1.** Properties of the 15 known disease-linked GOF mutants of KCNQ1.

## Acknowledgments

Flow Cytometry experiments were performed in the VMC Flow Cytometry Shared Resource and Veterans Affairs Tennessee Valley Healthcare System Flow Cytometry Core.

## Author contributions

HH, LM, and CGV conducted the experiments of this work. All authors participated in data analysis. CRS, HH, JPS, ALG, CGV, GK, and JM wrote the paper. CRS, ALG, and JPS conceived of this work and directed the approaches used.

