## Supplemental Figures and Table for "Disease-Linked Super-Trafficking of a Mutant Potassium Channel"

**Figure S1**

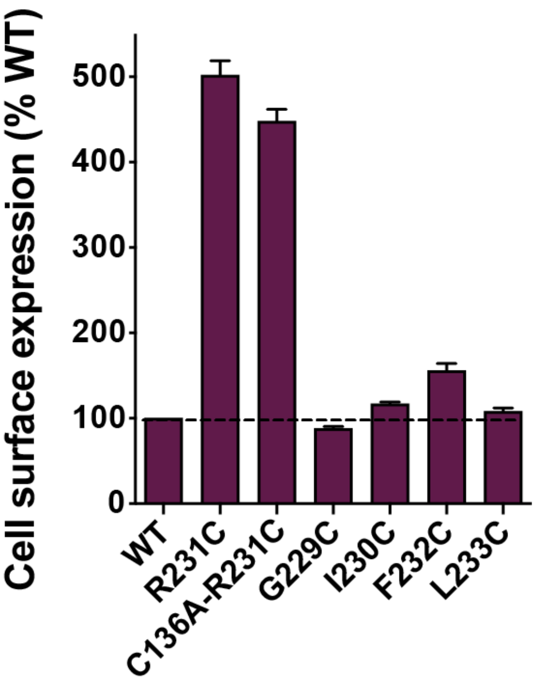


**Figure S1. Super-trafficking of R231C KCNQ1 is unrelated to disulfide bond formation.** Residue R231 is proximal to C136 in S1 when the channel is in the intermediate state, indicating that C231 could conceivably form a disulfide bond with C136. An Ala mutation is introduced to avoid any such potential disulfide bond. Residues surrounding R231 have been mutated to Cys to investigate whether mutation of any of site 231-proximal positions leads to super-trafficking. Bar height indicates mean value and the error bar is S.E.M. of at least three independent biological replicates.

**Figure S2**

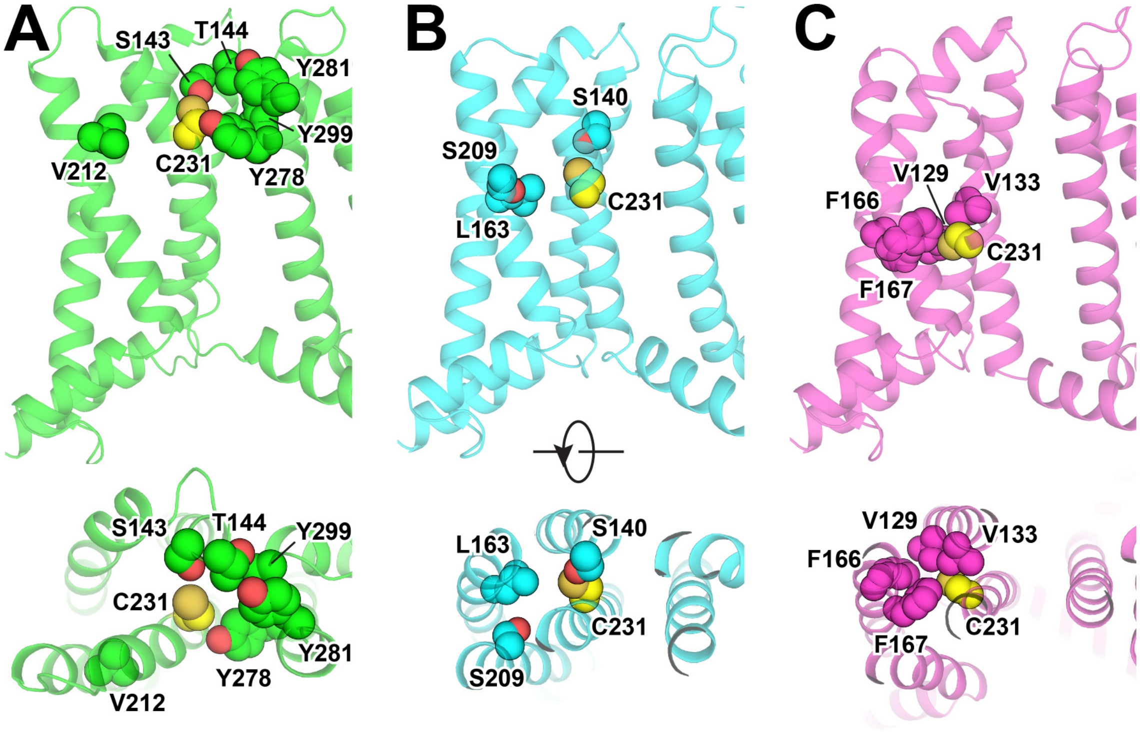


**Figure S2. Locations of residues used for double mutant analysis of interresidue side chain interactions involving C231 are illustrated in KCNQ1 structures**. Residues that are proximal to position 231 in current experimental and models structure of the activated, intermediate, and resting state of KCNQ1 are shown. (**A**) Human KCNQ1 activated state structure (PDB: 6V01) (*41*). (**B**) Rosetta model of the KCNQ1 intermediate state structure developed based on the experimental KCNQ1 VSD NMR structure (PDB: 6MIE) (*42*). (**C**) Rosetta model of the KCNQ1 resting state(*43*). The native R231 in these structural models was substituted for Cys, and the models were minimized with Rosetta. The side chains of C231 and of its proximal residues are depicted as spheres.

**Figure S3**

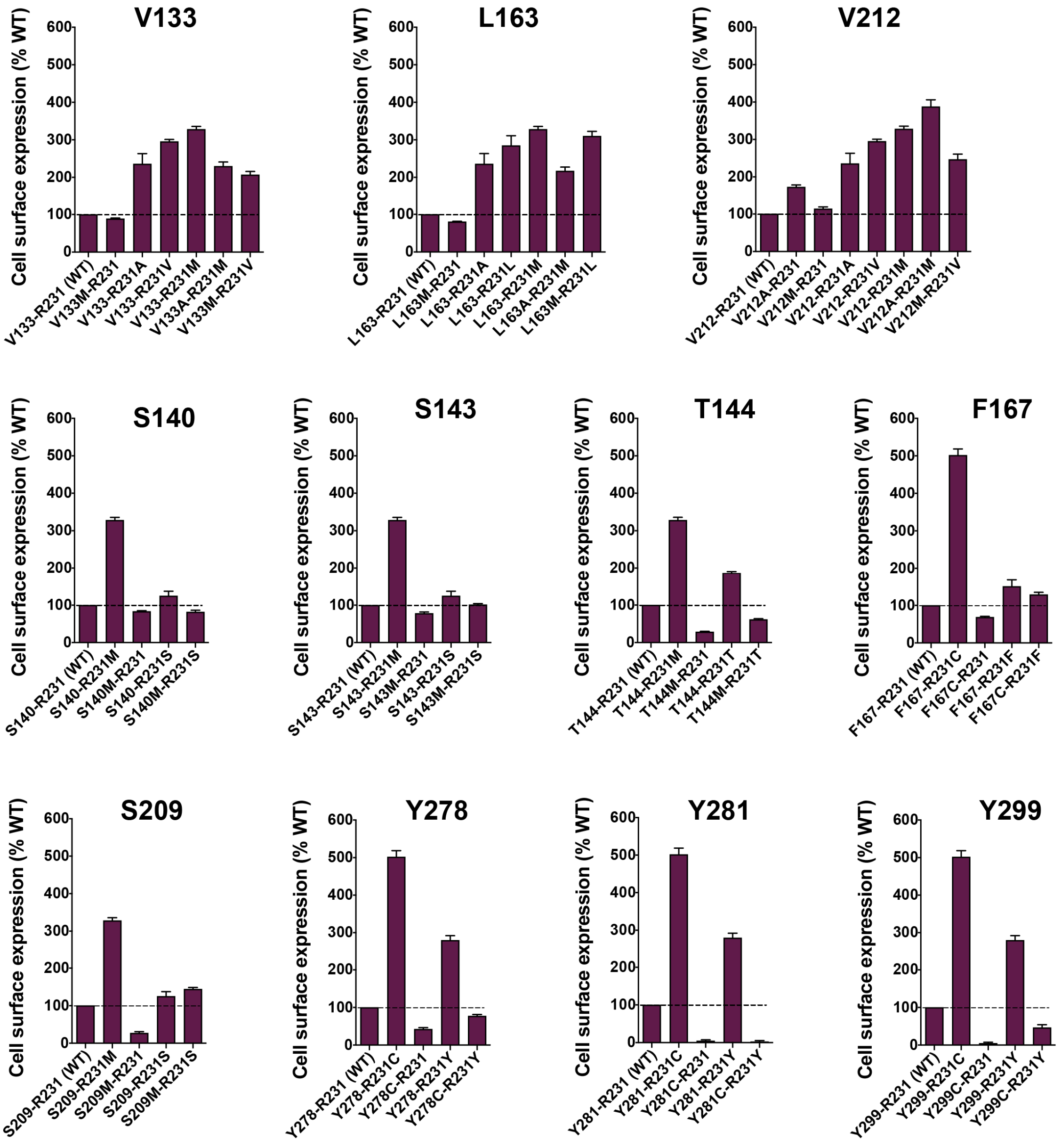


**Figure S3. Double mutant analysis has been performed for sites that were deemed not to be energetically coupled to the side chain for site 231 (see Figure 6A).** The mean cell surface expression levels of double mutants and corresponding single mutants have been plotted. The bar height indicates mean value and the error bar is S.E.M. of at least three independent biological replicates.

### Table S1. Properties of the 15 known disease-linked GOF mutants of KCNQ1.

| Protein Change | Codon Change | Location | Associated Phenotype | ClinVar Interpretation | Functional Property | Reference |
|---|---|---|---|---|---|---|
| **R14C** | CGC>TGC | N-terminus | AF | not provided | w/ E1: WT like; if cell swelling, $I_{max}$ increased, activation $\Delta V_{1/2}$ left shift, slow deactivation. | (*25*) |
| **A46T** | GCG>ACG | N-terminus | LQTS, AF | uncertain significance | w/o and w/ E1: $I_{max}$ increased or WT like, fast activation, activation $\Delta V_{1/2}$ WT like. | (*23, 27, 28*) |
| **S140G** | AGC>GGC | S1 | AF | Pathogenic | w/o E1: fast activation, slow deactivation.<br>w/ E1 or E2: $I_{max}$ drastically increased; activation $\Delta V_{1/2}$ left shift and instantaneous activation, slow deactivation.<br>w/ E3: WT like. | (*12, 13*) |
| **V141M** | GTG>ATG | S1 | SQTS, AF | pathogenic or likely pathogenic | w/o E1: WT like.<br>w/ E1: $I_{max}$ increased, activation $\Delta V_{1/2}$ left shift and instantaneous activation, slow deactivation. | (*12, 18*) |
| **Q147R** | CAG>CGG | S1-S2 linker | LQTS, AF | not provided | w/ E1: $I_{max}$ reduced, activation $\Delta V_{1/2}$ WT like.<br>w/ E2: $I_{max}$ increased, activation $\Delta V_{1/2}$ WT like.<br>w/o E1, w/ E3 or E4: WT like. | (*21*) |
| **R195W** | CGG>TGG | S2-S3 linker | LQTS, AF | uncertain significance | w/o and w/ E1: $I_{max}$ increased; fast activation and slow deactivation. | (*19, 24, 27*) |
| **S209P** | TCC>CCC | S3 | AF | Pathogenic | w/o E1: slow deactivation.<br>w/ E1: activation $\Delta V_{1/2}$ left shift and instantaneous activation, fast activation, slow deactivation. | (*14*) |
| **G229D** | GGC>GAC | S4 | LQTS, AF | Pathogenic | w/ E1: activation $\Delta V_{1/2}$ left shift and instantaneous activation, $I_{max}$ reduced, slow deactivation. | (*16, 29*) |
| **R231C** | CGC>TGC | S4 | LQTS, AF | Pathogenic | w/o E1: slow deactivation.<br>w/ E1: activation $\Delta V_{1/2}$ left shift and instantaneous activation, $I_{max}$ reduced. | (*10, 17, 45*) |
| **R231H** | CGC>CAC | S4 | LQTS, AF | Pathogenic | w/ E1: activation $\Delta V_{1/2}$ left shift and instantaneous activation.<br>w/ E3: $I_{max}$ reduced. | (*23, 54*) |
| **V241F** | GTC>TTC | S4 | LQTS, AF | uncertain significance | w/ E1: activation $\Delta V_{1/2}$ left shift and instantaneous activation, slow deactivation. | (*20*) |
| **I274V** | ATC>GTC | S5 | LQTS, SQTS, AF | conflicting interpretations | w/o E1: WT like.<br>w/ E1: $I_{max}$ increased, fast activation, slow deactivation. | (*26*) |
| **F279I** | TTT>ATT | S5 | SQTS | Pathogenic | w/o E: $I_{max}$ increased, slow activation, reduced inactivation.<br>w/ E1: activation $\Delta V_{1/2}$ left shift, fast activation, co-assembly with E1 decreased. | (*22*) |
| **V307L** | GTG>CTG/TT G | Pore helix | SQTS | Pathogenic | w/ E1: activation $\Delta V_{1/2}$ left shift and instantaneous activation, fast activation, slow deactivation. | (*11, 15*) |
| **R670K** | AGG>AAG | C-terminus | AF | not provided | w/o and w/ E1: $I_{max}$ increased. | (*24, 27*) |

LQTS: long QT syndrome  SQTS: short QT syndrome  AF: atrial fibrillation  E: KCNE
